# Notch Intracellular Domain Plasmid Delivery via Poly(lactic-co-glycolic acid) Nanoparticles to Upregulate Notch Signaling

**DOI:** 10.1101/2021.04.16.440241

**Authors:** Victoria L. Messerschmidt, Aneetta E. Kuriakose, Uday Chintapula, Samantha Laboy, Thuy Thi Dang Truong, LeNaiya A. Kydd, Justyn Jaworski, Kytai T. Nguyen, Juhyun Lee

## Abstract

Notch signaling is a highly conserved signaling system that is required for embryonic development and regeneration of organs. When the signal is lost, maldevelopment occurs and leads to a lethal state. Liposomes and retroviruses are most commonly used to deliver genetic material to cells. However, there are many drawbacks to these systems such as increased toxicity, nonspecific delivery, short half-life, and stability after formulation. We utilized the negatively charged and FDA approved polymer poly(lactic-co-glycolic acid) to encapsulate Notch Intracellular Domain-containing plasmid in nanoparticles. In this study, we show that primary human umbilical vein endothelial cells readily uptake the nanoparticles with and without specific antibody targets. We demonstrated that our nanoparticles also are nontoxic, stable over time, and compatible with blood. We also determined that we can successfully transfect primary human umbilical vein endothelial cells (HUVECs) with our nanoparticles in static and dynamic environments. Lastly, we elucidated that our transfection upregulates the downstream genes of Notch signaling, indicating that the payload was viable and successfully altered the genetic downstream effects.

## Introduction

Notch signaling is highly conversed cell signaling pathway which involved in diverse embryonic organs or tissue development as well as regeneration [1–10]. Notch signaling regulates cell-fate determination during activation by signal sending and receiving affected through ligand-receptor crosstalk. During the cell-fate decisions in cardiac[8, 11, 12], neuronal[13–15], immune[16, 17] and endocrine[18, 19] development, Notch signaling pathway is a key regulator including cell proliferation and differentiation [4, 20, 21]. Notch receptors are single-pass transmembrane proteins composed of functional Notch extracellular domain (NECD), transmembrane (TM), and Notch intracellular domains (NICD). Notch receptors are processed in the endoplasmic reticulum and Golgi apparatus within the signal-receiving cell through cleavage and glycosylation, generating a Ca^2+^-stabilized heterodimer composed of NECD non-covalently attached to the transmembrane NICD inserted in the membrane called S1 cleavage.

Regulation of arteriovenous specification and differentiation in both endothelial cells and vascular smooth muscle cells are also involved in Notch signaling including regulation of blood vessel sprouting, branching during normal and pathological angiogenesis, and the physiological responses of vascular smooth muscle cells [4, 6, 7, 22–24]. Defects in Notch signaling also cause inherited vascular diseases [4, 7, 23]. In endothelium, Delta-like ligand 4 (DLL4) is one of main ligand to send a signal to Notch in the adjacent cell [4, 6]. This in turn signals the surrounding cells to determine the cell-fate [4]. Once Notch is activated, the NICD is cleaved by γ-secretase and travels into the nucleus. Here, the NICD binds directly to the DNA, physically moving corepressors and histones, recruiting coactivators, and activating gene transcription [4, 6, 21].

When a disruption in this process occurs, either by chemical or genetic means, it causes developmental problems. For example, significant reduction in trabeculation is usually associated with deficient compaction in the ventricle [6, 25]. It has been shown that lack of trabeculation results in the inability to dissipate the kinetic energy, resulting in a malformed heart due to a decrease in Notch related signaling [25, 26]. Interestingly, when given NICD mRNA injection treatment, the heart function – including end diastolic function, end systolic function, stroke volume, and ejection fraction – were all partially or fully restored by rescuing downstream Notch signaling [27, 28]. There data demonstrated the possible impact of spatiotemporal NICD treatment for therapeutic approach to rescue Notch signaling.

Traditionally, retroviruses or liposomes have been used to deliver cDNA plasmids [2, 29–31]. These methods have various benefits such as DNA protection and DNA viability, but also have limitations of nonspecific delivery, stability after formulation, or host immune responses [32, 33]. Therefore, many groups are attempting to deliver the genetic materials such as cDNA plasmids via nanoparticles or nano-vehicles to mitigate these negative side effects. Various polymers have been used for gene delivery [34–39]. Cationic polymers have been used extensively to deliver genetic materials as DNA condenses quickly on the oppositely charges positive polymer. These polymers can be synthetic or organic and usually include polyethylenimine [40, 41], polyamidoamine [42, 43], chitosan [44, 45], and cationic proteins [46] or peptides. However, the drawbacks of these highly positively charged polymers are mainly due to its toxicity [32, 33] and often require extensive surface modifications to alleviate those effects [33]. Poly(lactic-co-glycolic acid) (PLGA), an FDA-approved biodegradable polymer [47], is a negatively charged polymer that has been extensively used for cancer treatment [48–51]. More recently, PLGA has been used to load both hydrophilic and hydrophobic materials such as cDNA plasmids [35] and RNAs [52], proteins [53–55], dyes [56], and drugs [57].

Therefore, we have here demonstrated that NICD DNA plasmid was successfully targeted and upregulate Notch signaling by using human umbilical vein endothelial cells (HUVECs) and flow chamber mimicking the circulation system.

## Methods

### Nanoparticle Synthesis and Conjugation

Poly(D, L-lactide-*co*-glycolic acid) nanoparticles (PLGA, 50:50, Akina Inc., West Lafayette, IN, USA) of two different molecular weights including 55–65 kDa (HMW Nanoparticles) and 5–10 kDa (LMW Nanoparticles) were fabricated by a standard double emulsion method as previously described [38]. In brief, PLGA was dissolved in chloroform (Sigma-Aldrich, St. Luis, MO, USA) at a 20 mg·mL^−1^ concentration. Following which, the water phase with 1% (w/w) rhodamine B was added to the oil phase dropwise under stirring and sonicated. The primary emulsion is then emulsified into 5% (w/v) Poly(vinyl) Alcohol (PVA, 13 kDa, Sigma) solution and then sonicated at 40 Watts for 5 minutes (30 sec off every 1 minute). Nanoparticles were then collected via centrifugation at 15,000 RPM for 15 minutes, then lyophilized until completely dry. Coumarin-6 loaded PLGA nanoparticles were prepared to track the nanoparticles’ interaction with the cells. For this, coumarin-6 was added into the oil phase at a ratio of 1:100 with respect to the amount of PLGA used during the nanoparticle synthesis. Rhodamine B loaded nanoparticles were exclusively used for dye release studies.

pCAG-GFP or TetO-FUW-NICD loaded HMW nanoparticles were also prepared based on the same standard double emulsion method with slight modifications according to past literature [58]. In brief, 250 μg of plasmid was diluted in 5% glucose solution to 200 μL which was then emulsified into 0.5 mL of 5% (w/v) PLGA solution in chloroform using a probe sonicator at 40W energy output for 15s to form primary water/oil emulsion. The primary emulsion was then emulsified into 3 mL of 4% (w/v) PVA solution by sonication and later dropped into 7.5 mL of 0.3% (w/v) PVA solution. The final mixture was then stirred for 3 hours at room temperature and particles were collected by centrifugation. Nanoparticles were then lyophilized until completely dry before use.

PLGA nanoparticles were conjugated either with anti-EGFL7 antibody (ab92939, Abcam) or anti-Tie2+1 antibody (ab151704, Abcam) via EDC-NHS chemistry as described elsewhere with modification [38]. In brief, nanoparticles were suspended in 0.1M MES buffer at a concentration of 2 mg/ml. Following which, 120 mg of EDC and 150 mg of NHS was added into the solution. After 2 hours of incubation at room temperature, nanoparticles were collected by centrifugation and resuspended in PBS (2mg/ml). 25μL of antibody solution was added into nanoparticles solution and incubated overnight at 4°C. The supernatant was used to determine the antibody conjugation efficiency using Bradford assay following manufacturers’ instructions. Pellets were resuspended in DI water, freeze-dried, and stored for use.

### Characterization and Stability of Nanoparticles

To determine the size and surface charge, nanoparticle suspension was added to a transparent cuvette and was then inserted into the ZetaPALS dynamic light scattering (DLS) detector (NanoBrook 90Plus PALS, Brookhaven Instruments, Holtsville, NY) as previously described [38]. Scanning electron microscopy (SEM, Hitachi S-3000N, Hitachi, Pleasanton, CA) was used to visualize the morphology of nanoparticles. Briefly, 50 μl of the nanoparticle suspension air-dried on a coverslip was silver sputter-coated and inserted into the SEM instrument. To determine the *in vitro* stability, nanoparticles were suspended in saline (0.9% Sodium Chloride, NaCl, Crystalline, Fisher Scientific, Hampton, NH, USA) or Vasculife VEGF basal cell media with 10% Fetal Bovine Serum (LL-0003, Lifeline Cell Technologies) and incubated at 37°C for 48 hours. Particle size was measured on predetermined time points using DLS as described earlier. The stability of the nanoparticles were represented as the percentage change of nanoparticle size measured at each time point with respect to initial particle size.

### Loading and Release Studies

The encapsulation efficiency of entrapped reagent including, pCAG-GFP or TetO-FUW-NICD, within PLGA nanoparticles was determined based on indirect loading analysis. Briefly, the un-loaded reagent in the supernatant (PVA solution) following the nanoparticle synthesis, was used to calculate the encapsulation efficiency (Equation 1). The amount of plasmid was determined using Picogreen DNA assay (#E2670, Promega, Madison, WI) following the manufacturers’ instructions.

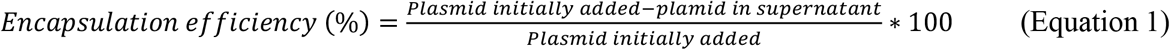

For *in vitro* plasmid release studies, solutions of pCAG-GFP or TetO-FUW-NICD plasmid-loaded nanoparticles were prepared in 1X PBS at a concentration of 1.5 mg/mL. At predetermined time points, the samples were centrifuged at 12,000 RPM for 5 minutes. The supernatant was then collected and stored at −20°C for further analysis. Pellet was again resuspended in fresh 1mL of PBS solution and incubated until next time point. Four replicates were used for analysis. For analysis, the plasmid solutions were incubated for Nb.Bsmi nicking enzyme (R0706S, New England Biolabs) for 60 minutes at 65°C in NEBuffer 3.1. The enzyme was then inactivated for 20 minutes at 80°C. The nicked plasmid supernatant was analyzed for plasmid release using the Picogreen DNA assays. The plasmid standards were made to determine the cumulative percentage of plasmid release over time.

### In vitro Compatibility of Nanoparticles

HUVECs were cultured in M199 media (M4530, Sigma-Aldrich) supplemented with Vasculife VEGF growth factor kit (LS-1020, Lifeline Cell Technologies) up to passage 7 in a 5% CO_2_ environment. To determine the compatibility of nanoparticles, HUVECs were seeded in 96 well plates at seeding density of 8,000 cells/well and cultured overnight. HMW nanoparticles and LMW nanoparticles of various concentrations (25, 50, 100, 250, 500, 1000 μg mL^−1^) were prepared in complete M199 media and added to the cells. After 24 hours of incubation at 37°C, the nanoparticle containing media was removed, and cells were carefully washed with 1X PBS. The cellular viability was then determined using MTS assays per manufacturer’s instructions.

In addition, HMW nanoparticles and LMW nanoparticles compatibility was evaluated using human whole blood, to determine hemolysis and whole blood clotting kinetics assay as previously mentioned. For these studies, whole blood was drawn from healthy adult volunteers into acid citrate dextrose anticoagulant tubes (ACD, Solution A; BD Franklin Lakes, NJ). Consent from the volunteers was obtained prior to the blood collection, and all the procedures strictly adhered to the IRB standards approved at the University of Texas at Arlington.

To perform whole blood clotting study, the blood was initially activated by adding 0.01 M of calcium chloride (Sigma). Following which, 50 μL of activated blood was added into 10 μL of saline diluted nanoparticle solution at concentration of 1 mg/mL and incubated for predetermined time points. At each time point, 1.5 mL of DI water was added to lysed the un-clotted blood and the absorbance of the supernatant was measured at 540 nm. Untreated blood served as a control. In hemolysis study, nanoparticles were suspended in saline at the following concentrations (0, 10, 25, 50, 100, 250, 500, 1000 μg·mL^−1^) and then incubated with 200 μL of saline-diluted blood for 2 hours at 37°C. Following the incubation, the samples were centrifuged and the absorbance of the supernatant was quantified at 545 nm. Untreated blood that was diluted with DI water and saline solution served as positive and negative controls respectively. The percent hemolysis was calculated using the following equation:

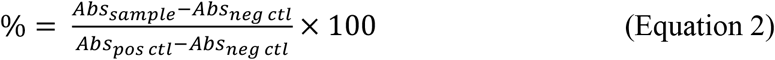

### In vitro Cellular Uptake and Interaction of Nanoparticles

To determine the uptake of coumarin-6 loaded HMW- and LMW-PLGA nanoparticles by HUVECs, cells were seeded in 96 well plates at a density of 8,000 cells/well. After overnight culture, nanoparticles of various concentrations 50, 100, 250, 500, 1000 μg·mL^−1^ were added to the cells and incubated for 4 hours in 37°C. Nanoparticles were then removed, cells were carefully washed with PBS solution and lysed using 1% Triton X-100. Fluorescence intensity measurement of nanoparticles in cellular lysate was quantified at a wavelength of 457 nm (excitation)/500 nm (emission) using a spectrophotometer. These measurements were analyzed against a nanoparticle standard. The measurements were further normalized with respect to the sample cellular protein amount as determined based on BCA assays (Thermofisher Scientific).

Similarly, interaction between antibody (anti-EGFL7 or anti-Tie2&Tie1) conjugated HMW nanoparticles loaded with coumarin-6 and HUVECs were also determined under static conditions. In brief, nanoparticle suspensions were treated with cells for 30 minutes and following which, cells were washed and lysed. Cellular lysate was used to determine the amount of nanoparticle attachment and internalization with HUVECs based on coumarin-6 fluorescence intensity. These fluorescence measurements values were then normalized with the total DNA content per sample using Picogreen DNA assays per manufacturer’s instructions. In parallel, nanoparticle interaction with HUVECs were observed using a fluorescence microscope under FITC channel. The cells were counterstained using Nucblue (Invitrogen) to visualize the cell nuclei.

In addition, the ability of a coumarin-6 loaded, antibody (anti-EGFL7 or anti-Tie2&Tie1) conjugated HMW nanoparticles to adhere and interact with HUVECs under physiological relevant flow condition was investigated. HUVEC’s were seeded at 2*10^6^ cells/mL into μSlide VI^0.4^ channel and cultured overnight. Following the cell attachment, nanoparticles suspended in M199 media at a concentration of 200 μg/mL were perfused through the channels of the flow slide using Ibidi pump system at a shear stress of 5 dyne/cm^2^ for 30 minutes. Later, cells within the channels were fixed with paraformaldehyde solution and treated with Nucblue (Invitrogen) to stain cell nuclei. The cellular images were then taken using fluorescence microscope under FITC and DAPI channel to visualize the nanoparticles and nuclei, respectively. The fluorescence intensity of nanoparticles was later quantified using NIH ImageJ software and normalized by cell number.

### Plasmid Transfection

HUVECs were seeded 24 hours prior to the transfection study at n=4. The following day, Lipofectamine 3000 or no treatment were applied to the cells for 6 hours. After the treatment, the cells were washed three times with 1X PBS and incubated until the next time point. HMW PLGA nanoparticles were prepared as described above. The nanoparticles at a concentration of 250 μg/mL were then applied to HUVECs for 6 hours. The cells were then gently washed with 1X PBS three times and new media given. The cells treated with Lipofectamine, nanoparticles, or no treatment were then grown for 24, 48, or 72 hours post transfection. Cells transfected with pCAG-GFP plasmid-loaded nanoparticles were imaged in a fluorescent microscope on FITC channel. Those that were subjected to TetO-FUW-NICD plasmid-loaded nanoparticles were analyzed via RT-PCR.

### RT-PCR Data

Cells were first washed with 1X PBS two to three times. Then, 0.025% trypsin was added for 3 minutes at 37°C to allow cell detachment. The trypsin was then neutralized by adding media twice the volume of trypsin to the wells. The cells were collected, centrifuged at 150×g for 5 minutes, and the supernatant discarded. The cells were then used to isolate the total RNA using the Aurum Total RNA Mini Kit (Biorad, #7326820) following the manufacturer’s instructions. RNA concentration was determined via NanoDrop, by reading each sample 3 times. The total RNA was then used to synthesize 200 ng of cDNA using the iScript Synthesis Kit (Biorad, #1708890) following the manufacturer’s instructions. PCR was conducted using the iTaq Universal SYBR Green Supermix (Biorad, #1725121) following manufacturer’s instructions. The primer sequences for human mRNA are as follows: *Dll4* (Frd CTGCGAGAAGAAAGTGGACAGG, Rev ACAGTCGCTGACGTGGAGTTCA), *Hes1* (Frd GGAAATGACAGTGAAGCACCTCC, GAAGCGGGTCACCTCGTTCATG*), Hey1* (Frd ACCATCGAGGTGGAGAAGGA, Rev AAAAGCACTGGGTACCAGCC), *Notch1* Receptor (Frd GGTGAACTGCTCTGAGGAGATC, Rev GGATTGCAGTCGTCCACGTTGA), NICD (Frd ACCAATACAACCCTCTGCGG, Rev GGCCCTGGTAGCTCATCATC), and *β*-*Actin* (CGACAGGATGCAGAAGGAG, Rev ACATCTGCTGGAAGGTGGA).

## Results

### Optimization of Nanoparticles based on Molecular Weight of PLGA

Before performing the cell study, nanoparticles were characterized based on their size, poly dispersity, and zeta potential (**Table 1**). Among the two molecular weights tested, the high molecular weight (HMW) PLGA nanoparticles, at 55–65 kDa, were smaller than the low molecular weight (LMW), 1–5 kDa PLGA, nanoparticles at 234 ± 90 and 246 ± 85 nm, respectively. The zeta potential, or surface charge of the nanoparticles, indicates the presence of the negatively charged carboxyl and hydroxyl groups present on the polymer. The HMW PLGA nanoparticles have a charge of −31 ± 3.4 mV, and the LMW PLGA nanoparticles have a charge of −29 ± 2.8 mV. The poly dispersity of both the HMW and LMW PLGA nanoparticles, 0.13 ± 0.05 and 0.08 ± 0.02 respectively, shows that the particles are uniformly dispersed. SEM images also indicated that both the HMW- and LMW-nanoparticles were uniformly dispersed and have spherical morphology (**Figure 1A**).

**Table 1:** PLGA Nanoparticle Physical Attributes.

| PLGA Nanoparticles | Size (nm) | Poly Dispersity | Zeta Potential (mV) |
| --- | --- | --- | --- |
| **MW: 55 – 65 kDa** | 234 ± 90 | 0.13 ± 0.05 | -31 ± 3.4 |
| **MW: 1 – 5 kDa** | 246 ± 85 | 0.08 ± 0.02 | -29 ± 2.8 |

**Figure 1:**
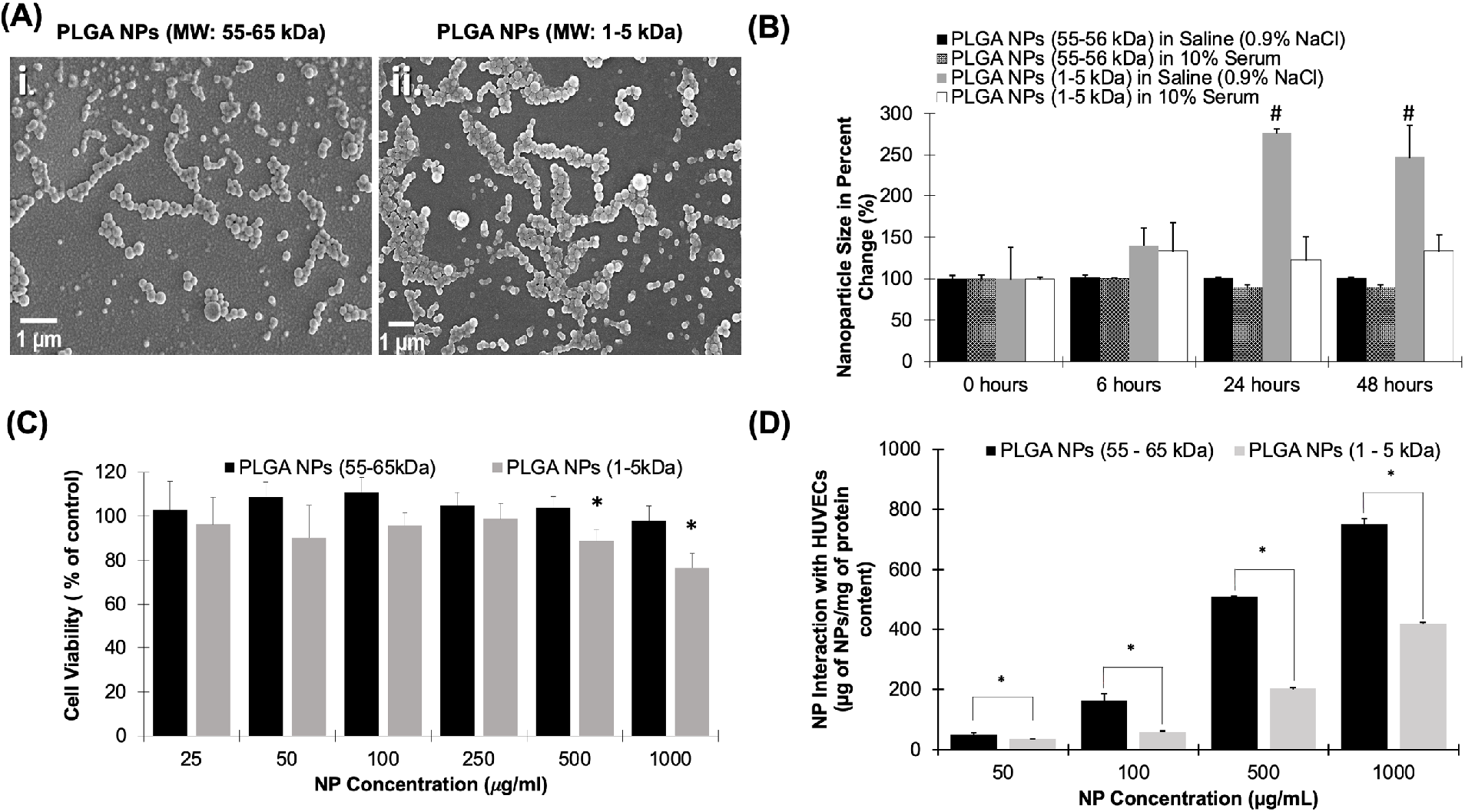
Characterization of PLGA Nanoparticles. *(A) SEM images of (i) PLGA at 55 – 65 kDa and (ii) PLGA at 1– 5 kDa nanoparticles. Scale bar is 1 μm. (B) Nanoparticles stability in saline (0.9% NaCl) or 10% Serum over time shows that the HMW PLGA nanoparticles’ size is steady in both solutions, while the LMW PLGA nanoparticles vary in both the Saline and 10% Serum. Error bars denote standard error. # indicates a significant difference from the samples’ time point 0 (p < 0.05). (C) Cytocompatibility test comparing HMW to LMW PLGA nanoparticles. This shows at all concentrations the HMW PLGA nanoparticles are insignificantly toxic to the cells. The LMW nanoparticles are tolerated up to a concentration of 250 μg/mL. * indicates a significant difference from 100% viability. (D) HUVEC uptake of both HMW and LMW PLGA nanoparticles shows HMW had significantly higher uptake than LMW at all concentrations. Data shown as mean + standard error. * indicates a significant difference between HMW and LMW Nanoparticle uptake*.

Following *in vitro* stability studies using HMW- and LMW-nanoparticles in both saline (0.9% NaCl) and 10% serum, the nanoparticle percent size change was determined. Accordingly, the diameter of HMW nanoparticles were constant in both formulations over 48 hours of incubation, which indicates the superior stability properties of HMW nanoparticles. On other hand, the size of LMW nanoparticles steadily increased over time and showed significant aggregation following their incubation with the saline solution at 24 hours. In serum, the LMW nanoparticles increased in size, but was not significantly different (**Figure 1B**). This suggests that LMW nanoparticles were not stable and may exhibit aggregation behavior following their suspension and/or administration. Then, the drug release kinetics were then compared between the two molecular weights. Using rhodamine B as a model hydrophilic drug, high and low molecular weight nanoparticles were incubated in 1X PBS over a period of time. After 9 hours, the LMW nanoparticles were able to release 48.5% of the loaded dye, compared to the 8.4% release of the HMW nanoparticles. After 5 days incubation, the LMW nanoparticles had released 100% of the Rhodamine B, while the HMW nanoparticles reached 20% of loaded dye (**Figure S1**). Both molecular weights of PLGA nanoparticles showed a burst release of rhodamine B dye, and the HMW nanoparticles had a sustained release until day 28 (**Figure S1**).

To assess the cytocompatibility of nanoparticles, HUVECs were subjected to varying concentrations of both HMW and LMW nanoparticles. Across all tested concentrations, the HMW nanoparticles were all above 90% viability, while the LMW had greater than 90% viability in only 25, 50, 100, and 250 μg/mL **(Figure 1C)**. At both 500 and 1,000 μg/mL, the LMW nanoparticles were significantly lower at 88 ± 10% and 76 ± 13% viability, respectively (p < 0.05). The uptake of the nanoparticles was evaluated using HUVECs incubated with varying amounts of nanoparticles. At each tested concentration, the HMW nanoparticles had a significantly higher uptake compared to that of the LMW. Additionally, there is a trend showing a dose-dependent relationship between the number of nanoparticles applied, and the number of nanoparticles endocytosed by the cells (**Figure 1D**).

### Hemocompatibility

To simulate the effect of nanoparticles on human blood, hemolysis and whole blood clotting tests were conducted. For whole blood clotting, the nanoparticles significantly affected the clotting cascade only during the first 10 minutes of exposure. Afterwards, the progress of blood clotting gradually reduced and there was no significant results compared to whole blood without exposing PLGA nanoparticles (Figure 2A). After 60 minutes, blood exposed to either HMW and LMW nanoparticles had great low supernatant absorbance, 0.1, similar to whole blood which indicates blood clot. Our results reflect those who have performed similar studies showing little red blood cell lysis or reduced clotting kinetics [59]. Furthermore, the interaction between red blood cells and nanoparticles were evaluated by incubation with diluted blood to determine if hemolysis occurred. Compared to lysed cells as the positive control, both the HMW and LMW nanoparticles were significantly lower (< 0.25%) in hemolysis **(Figure 2B).**

**Figure 2:**
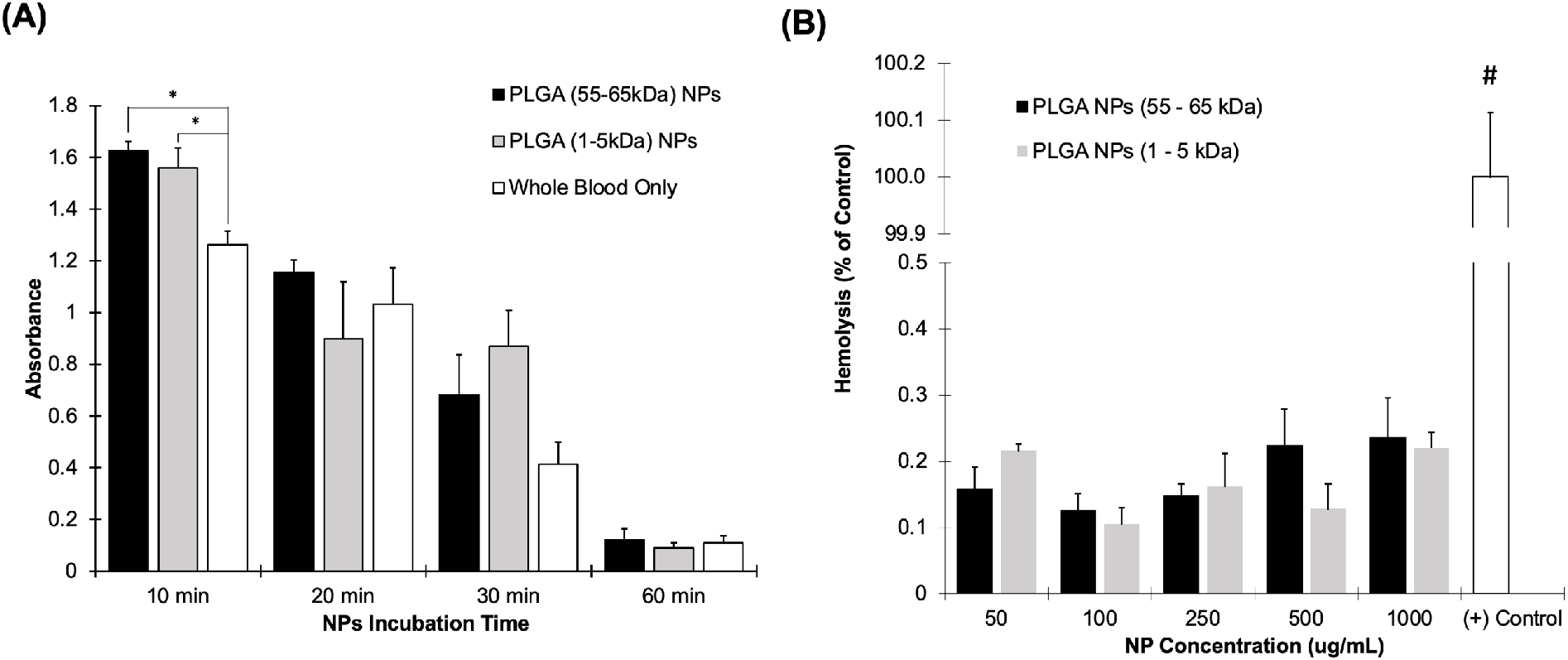
Hemocompatibility of PLGA Nanoparticles. *(A) Nanoparticles at a concentration of 1 mg/mL were subjected to human blood for up to 1 hour. The clotting was significantly affected only during the first 10 minutes of nanoparticle exposure (p < 0.05). At all other time points, the nanoparticles did not affect the clotting ability of the human blood. Blood exposed to only air was kept as a control. * denoted a significant difference with p < 0.05 (B) Nanoparticles were incubated with blood for 1 hour. Compared to RO water treatment (Positive control), both nanoparticle groups had significantly less hemolysis at all tested concentrations. # indicates that the Positive control is significantly higher than all other groups with p < 0.05. All data is shown as mean + standard error*.

### Selection of Optimal Endothelial Target

For this study, HMW nanoparticles were used due to the fact that the HMW has greater cell uptake and cell viability properties even though LMW nanoparticles have a rapid release profile. Anti-EGFL7 and Anti-Tie2+1 were conjugated to PLGA HMW nanoparticles and characterized. The nanoparticles conjugated with anti-EGFL7 became 249 ± 55 nm, while the nanoparticles conjugated with anti-Tie2+Tie1 are 243 ± 41 nm. Both antibody conjugations had a low poly dispersity, indicating that most of the nanoparticles were uniform in size. The antibodies changed the surface charge of the nanoparticles from −31 ± 3.4 to −23.5 ± 1.7 mV for anti-EGFL7 nanoparticles, and −31 ± 3.4 to −27.4 ± 1.8 mV for anti-Tie2+Tie1 nanoparticles. The antibodies had a conjugation efficiency of 59.6 ± 1.5% and 47.5 ± 1.2% for anti-EGFL7 conjugated nanoparticles and anti-Tie2+Tie1 conjugated nanoparticles, respectively (**Table 2**).

**Table 2:** Endothelial Cell Targeted PLGA Nanoparticles.

| Antibody Conjugated Nanoparticles | Size (nm) | Poly Dispersity | Zeta Potential (mV) | Conjugation Efficiency (%) |
| --- | --- | --- | --- | --- |
| **Anti-EGFL7 Nanoparticles** | 249 ± 55 | 0.21 ± 0.01 | -23.5 ± 1.7 | 59.6 ± 1.5 |
| **Anti-Tie2+Tie1 Nanoparticles** | 243 ± 41 | 0.19 ± 0.13 | -27.4 ± 1.8 | 47.5 ± 1.2 |

Furthermore, we tested our antibody conjugated particles on their binding abilities under static and physiological flow conditions. Under static culture, we saw concentration-dependent uptake of nanoparticles by endothelial cells (**Figure 3A**). Additionally, antibody conjugated nanoparticles had a greater interaction with the cells compared to unconjugated ones. Coumarin-6 loaded HMW nanoparticles conjugated to either anti-EGFL7 or anti-Tie2+Tie1 supported the quantitative data (**Figure S2**). Nanoparticles conjugated with anti-EGFL7 had significantly higher cellular uptake than unconjugated nanoparticles at a concentration of 100 μg/mL. Although anti-EGFL7 conjugated nanoparticles had higher cellular uptake than unconjugated nanoparticles at the other tested concentrations, they did not reach significance. On the other hand, nanoparticles conjugated with anti-Tie2+Tie1 were significantly higher in cellular uptake at all tested concentrations. As the concentration of anti-Tie2+Tie1 conjugated nanoparticles increases, the rate of cellular uptake increases 3.5-folds and 8.4 folds from 100 μg/mL to 250 and 500 μg/mL, respectively. Similarly, anti-EGFL7 conjugated nanoparticles increase 2.4-folds and 5.1-folds from 100 μg/mL to 250 and 500 μg/mL, respectively. Yet, the unconjugated nanoparticles increase 5.2-folds and 7.3-fold from concentrations of 100 to 250, and 500 μg/mL, respectively (**Figure 3A)**.

**Figure 3:**
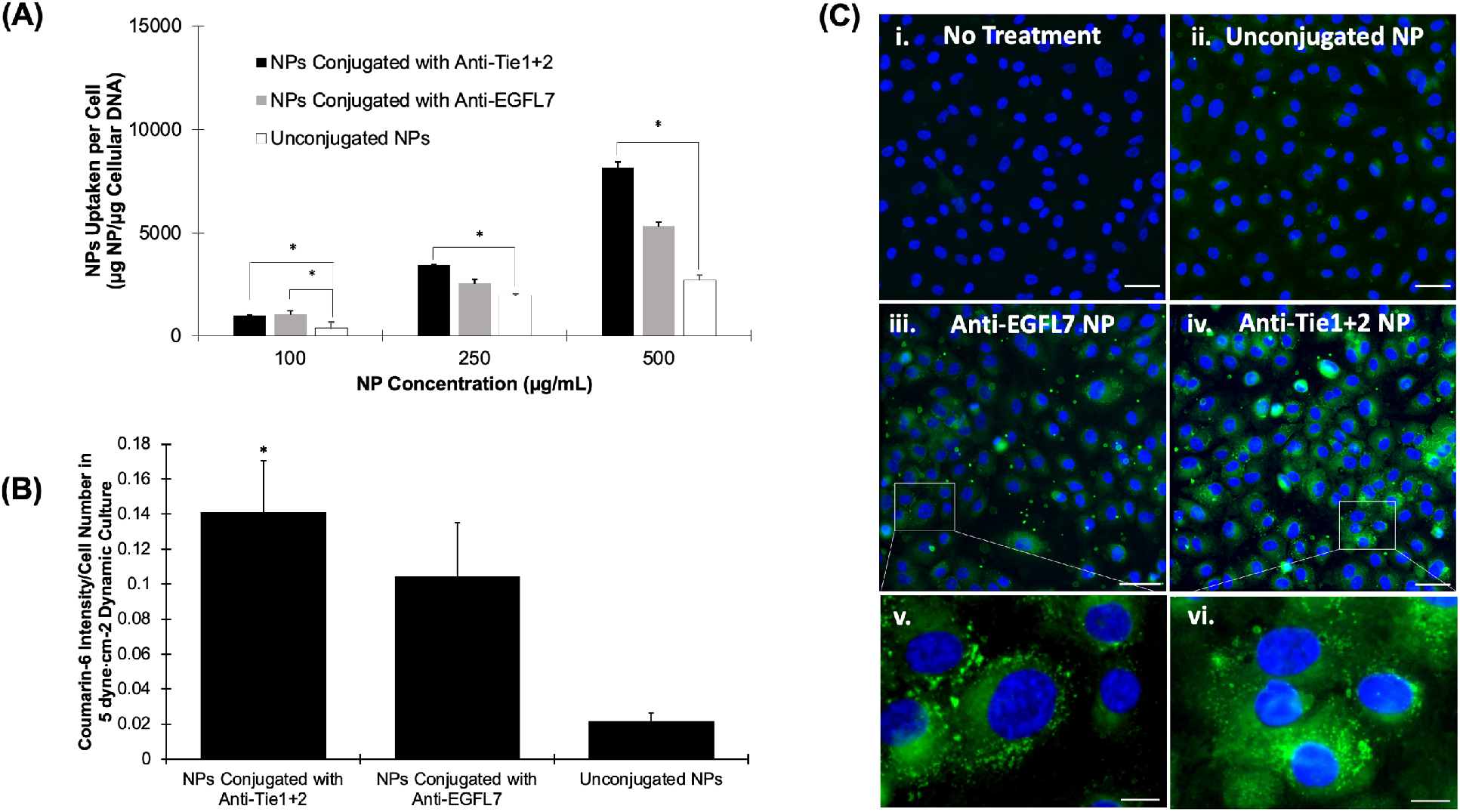
Targeting Efficiency of Antibody Conjugated PLGA Nanoparticles. *(A) HUVEC Cellular uptake of endothelium specific anti-EGFL7, anti-Tie1+2 conjugated nanoparticles or unconjugated nanoparticles after 4 hours. (B) HUVEC Cellular uptake under 5 dyn/cm^2 of endothelium specific anti-EGFL7 and anti-Tie1+2 conjugated nanoparticles, or unconjugated nanoparticles. (C) Fluorescent images of HUVEC’s incubated in flow culture of 5 dyn/cm^2 with (i) No Treatment, (ii) Unconjugated Nanoparticles, (iii) Anti-EGFL7 conjugated nanoparticles, (iv) anti-Tie1+2 conjugated nanoparticles. Scale bar = 20 um. Zoomed in portions of HUVEC’s incubated in flow culture at 5 dyn/cm^2 with (v) Anti-EGFL7 conjugated nanoparticles, and (vi) anti-Tie2+Tie1 conjugated nanoparticles. Scale bar = 5 um*.

In concordance with our observation in static condition, anti-Tie2+Tie1 conjugated nanoparticles were uptaken significantly higher than anti-EGFL7 conjugated nanoparticles at a concentration of 250 μg/mL under 5 dyne/cm^2^ (**Figure 3B**). Although both 250 and 500 μg/mL concentration of nanoparticles showed positive cell interaction, cell viability, and hemocompatibility, the concentration of 250 μg/mL was chosen for due to the higher downstream genetic effects and avoiding a potential negative feedback effect. In **Figure 3C**, we can see that under flow conditions, the antibody conjugated nanoparticles were able to be uptaken into cells at a higher rate compared to unconjugated nanoparticles. Due to the increase in cellular uptake of nanoparticles conjugated with anti-Tie2+Tie1, this antibody was determined to be superior for endothelial targeting.

### Characterization of Plasmid Loaded PLGA Nanoparticles

Both pCAG-GFP and TetO-FUW-NICD were loaded into HMW PLGA nanoparticles at 62.3 ± 2.2 μg plasmid per mg of nanoparticles and 89.1 ± 6.4 μg plasmid/mg nanoparticles, respectively. The encapsulation efficiency of 56.3 ± 4.1% and 38.9 ± 2.17% for NICD and GFP plasmids, respectfully, is similar to previous report [60–62] (**Table 3**). Additionally, previously reported particles were larger[61] and the encapsulated plasmids were 6- to 2-times smaller in size [60–62] of our largest plasmid, at 10,671 bp, the genetic material encapsulated into the nanoparticles was released in a similar form to our model drug, rhodamine B (**Figure 4A, S1, S3**). The NICD plasmid released up to 1 μg of plasmid within the first 24 hours. The plasmid continued to be released over 21 days to a total of 1.2 μg (**Figure 4A**). Our GFP plasmid loaded nanoparticles similarly released 0.5 μg of plasmid over 21 days (**Figure S3**). With the addition of the plasmids, the zeta potential and size both increased, indicating a change. However, the polydispersity value was still low illustrating their homogeneous size.

**Table 3:** NICD Plasmid Loaded PLGA Nanoparticle Characteristics.

|  | Size (nm) | Poly Dispersity | Zeta Potential (mV) | Encapsulated Efficiency (%) |
| --- | --- | --- | --- | --- |
| **NICD-Loaded PLGA Nanoparticle** | 272 ± 51 | 0.12 ± 0.05 | -12.9 ± 1.90 | 56.3 ± 4.1% |
| **NICD-Loaded PLGA Nanoparticle Conjugated with Anti-Tie2+Tie1** | 268 ± 26 | 0.11 ± 0.01 | -17.0 ± 0.83 |  |

**Figure 4:**
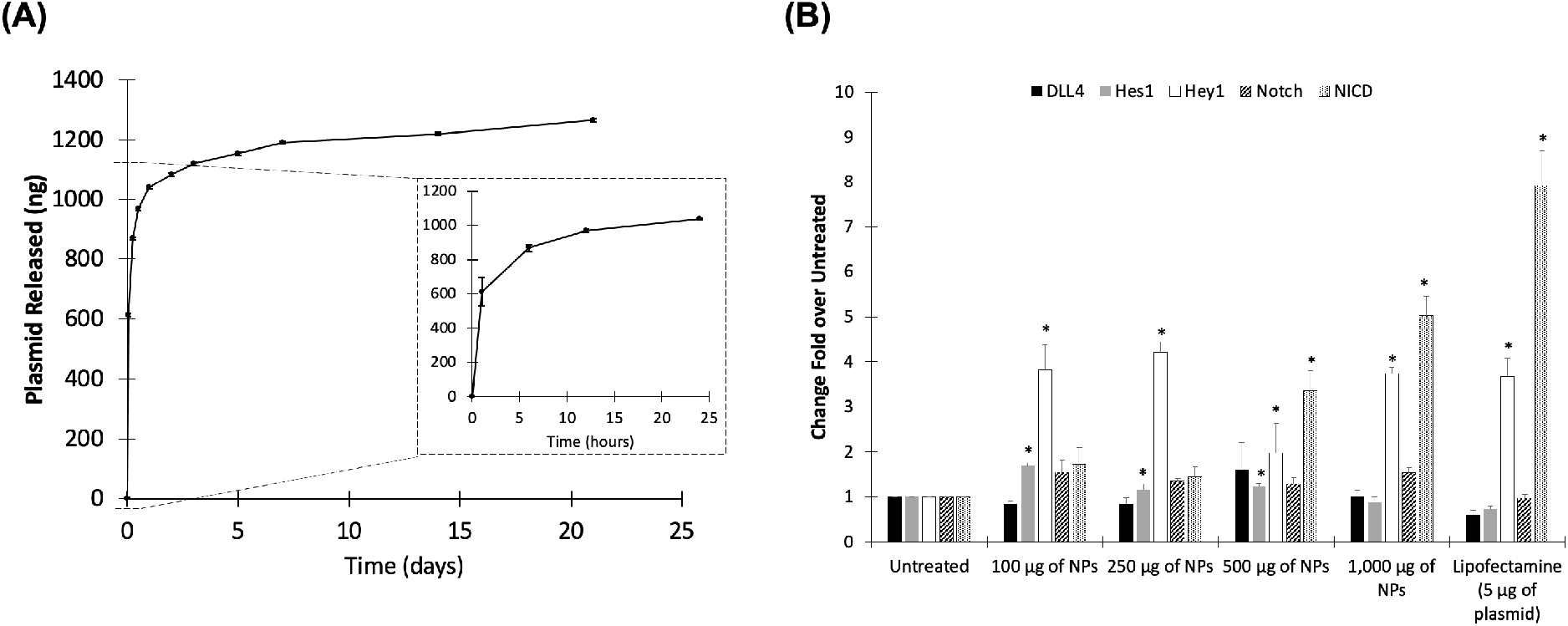
Characterization of NICD Loaded Nanoparticles. *Release curve of (A) NICD Plasmid-loaded nanoparticles measured via Promega dsDNA assay after 21 days. n = 3. Data shown as mean +/- standard deviation. (B) Quantitative expression of NICD after TetO-FUW-NICD Nanoparticle transfection at varying dosages. *significantly different from Untreated Group*.

TetO-FUW-NICD loaded nanoparticles were also given to HUVECs at varying doses. Compared to the untreated group, Notch target gene, *Hey1* was upregulated in each tested concentration. Additionally, *Hes1* was upregulated with NICD plasmid concentrations of 100, 250 and 500 μg of nanoparticles while 1000 μg of NICD plasmid loaded nanoparticle decrease the expression level of *Hey1* (**Figure 4B**). According to the NICD plasmid concentration of nanoparticle, however, expression level of NICD expectedly increase after treating to cells. Therefore, expression level of NICD does not proportion to Notch target gene expression.

### GFP Expression Over Time

HUVECs were subjected to 5 μg of plasmid, either through Lipofectamine 3000, (ThermoFisher), or our GFP Plasmid-loaded nanoparticles. After 6 hours, the treatments were removed, and fresh media applied to the cells. At 12 hours, there were not many cells that expressed GFP. At 24 hours, the lipofectamine group had much higher cell death than that of the GFP Plasmid nanoparticle group. Additionally, the GFP plasmid-loaded nanoparticles had an even expression of GFP across most cells. At 24 hours, the lipofectamine group had very few GFP positive cells compared to that of the nanoparticle treated group. At 48 hours post transfection, GFP was observed in both lipofectamine treated and GFP plasmid-loaded nanoparticle treated groups. The lipofectamine had a brighter GFP signal at 48 hours compared to the nanoparticle treatment, however, the nanoparticles transfected a higher number of cells, which lead the 5 μg of plasmid to be more evenly spread among cells (**Figure 5)**. The GFP plasmid-loaded nanoparticle group had a higher survival rate and high transfection efficiency.

**Figure 5:**
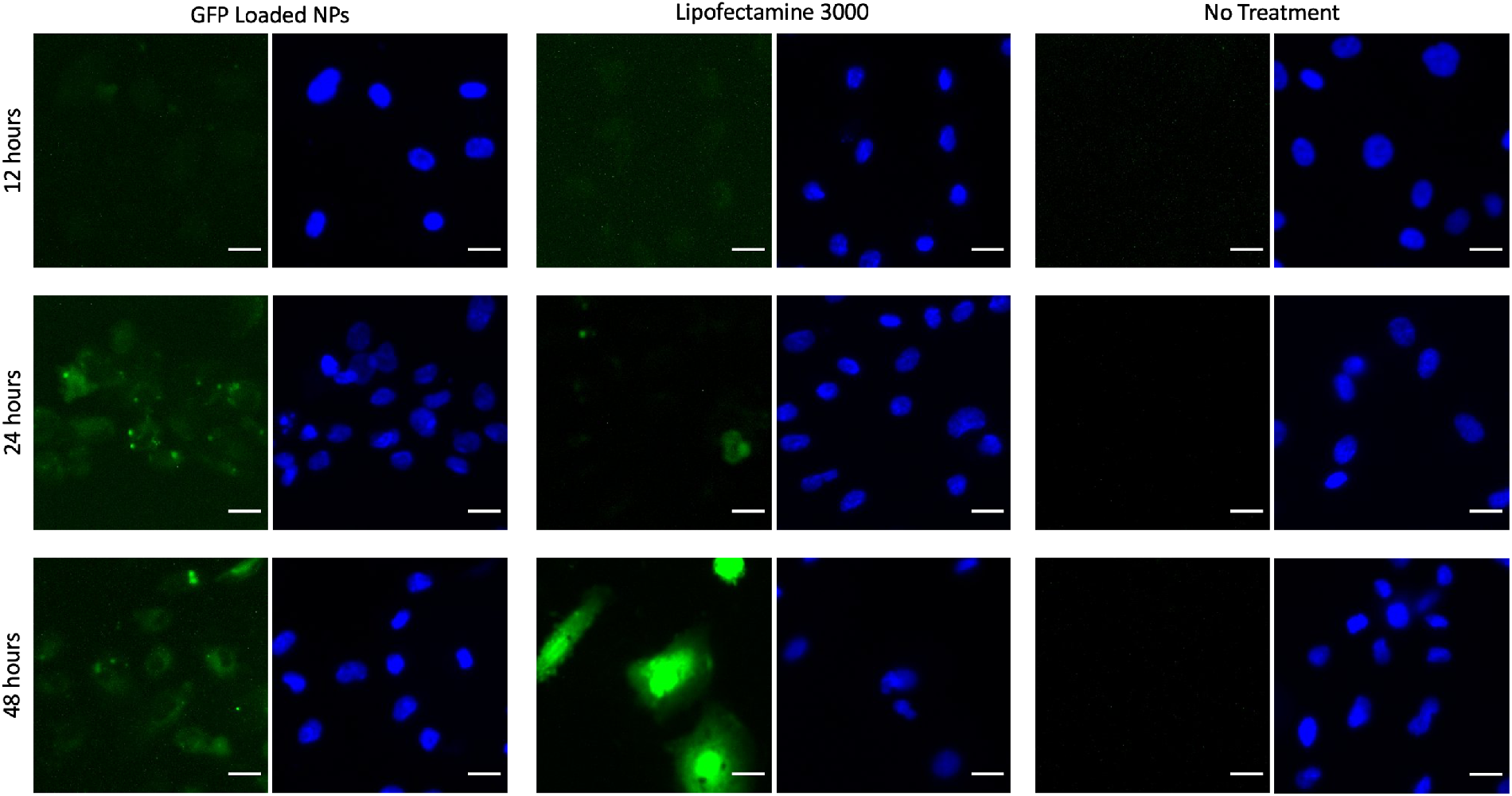
GFP Plasmid-loaded Nanoparticle Transfection. *After 6 hours of treatment, nanoparticles or lipofectamine were washed off with 1X PBS and media replaced. At 12 hours, slight GFP expression can be seen. After 24 hours, there is significantly more GFP expression in GFP plasmid-loaded nanoparticles than lipofectamine treated cells. Additionally, after 48 hours, lipofectamine transfected cells had a high GFP signal in few cells. Whereas nanoparticle transfected cells had many cells expressing GFP, resulting in a lower over signal*.

### Nanoparticle Mediated HUVEC transfection based on NICD Expression

HUVECs were subjected to 12 dyne/cm^2^ for 24 hours, then an additional 24 hours of flow with 2 μg/mL of doxycycline and treated with either blank nanoparticles, TetO-FUW-NICD loaded HMW nanoparticles, TetO-FUW-NICD loaded HMW nanoparticles conjugated with anti-Tie2+Tie1, or cell media only. Each nanoparticle group was given at a concentration of 250 μg/mL. The plasmid-loaded nanoparticles with targeting antibody had significantly higher expression of Notch related genes **(Figure 6)**. The expression of Notch related genes when exposed to plasmid-loaded nanoparticles without a conjugating antibody were not significantly different from that of the blank nanoparticles. Both were upregulated most likely due to the increased viscosity of the media due to the addition of the nanoparticles. The higher viscosity causes a higher shear stress, which Notch has been shown to respond to **(Figure S4)**.

**Figure 6:**
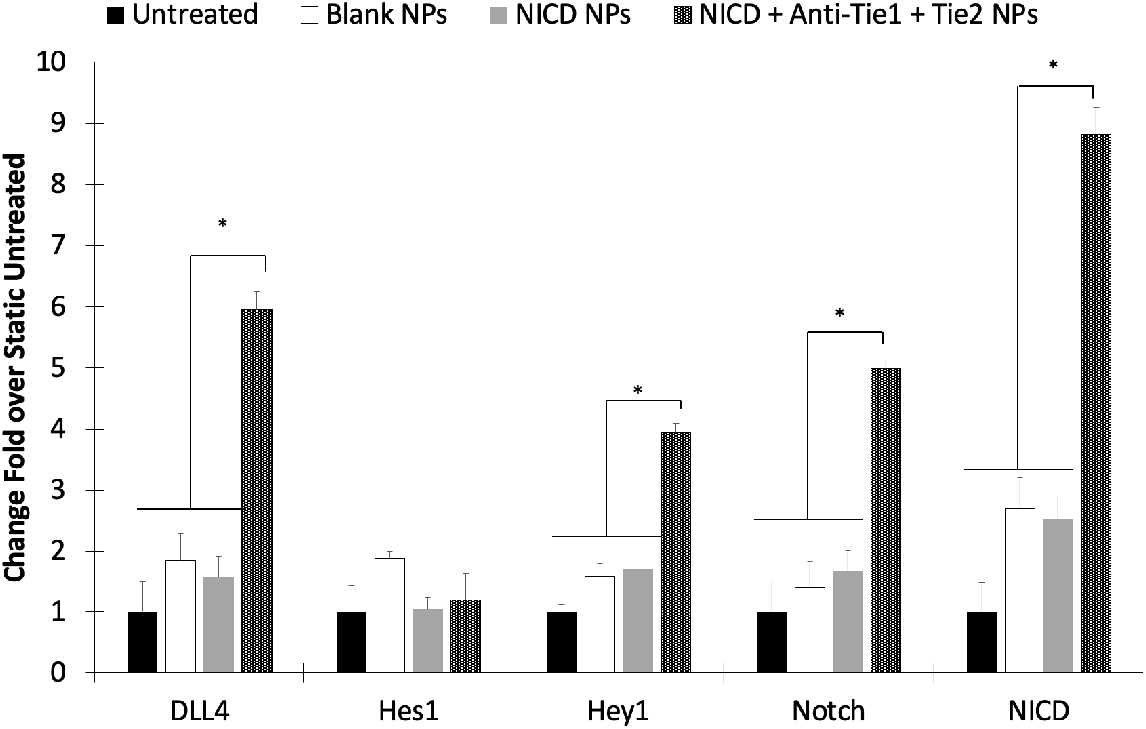
NICD Plasmid-loaded Nanoparticles can Enhance NICD Expression in Dynamic Culture Conditions. *RT-PCR results showing that NPs conjugated to anti-Tie2+Tie1 significantly upregulate DLL4, Hey1, Notch Receptor and NICD in 12 dyne/cm2 flow conditions. * NICD+Anti-Tie1+Tie2 is significantly higher (p < 0.05)*.

## Discussion

NICD as a transcription factor that is cleaved from Notch ligand and translocates into nucleus to activate Notch signaling which involve in cell proliferation and differentiation during development or tissue regeneration. Therefore, regulation of spatiotemporal activation of Notch is an important topic in developmental biology and regenerative medicine. In this paper, we have demonstrated the PLGA nanoparticle conjugated with NICD plasmid successfully upregulate Notch signaling.

Poly(lactic-co-glycolic acid) is one of the most characterized biopolymers with respect to drug delivery design and performance [63]. PLGA has been widely utilized for delivering proteins[53, 64–66] and hydrophobic drugs[67–70]. More recently, gene delivery for vaccines[34], immunotherapy [31], or gene therapy[40, 60, 62, 71, 72] have utilized nanoparticles. Traditionally, many studies have been used for viral or liposomal gene delivery for various applications[29, 71], but poses negative immunological effects, random gene integration, base pair size restrictions, and cytotoxicity [73]. As an FDA approved material, PLGA degrades via hydrolysis into glycolic and lactic acids, both of which occur naturally in the body [47, 62, 69]. Therefore, we used PLGA nanoparticles as a method of delivering genetic material into targeted cells. First, we optimized the molecular weight of the PLGA. Our data shows that the higher molecular weight PLGA was more cytocompatible, hemocompatible, and stable in various solutions. Even though the low molecular weight released the plasmid quickly, the nanoparticles were unstable in saline, an important liquid vehicle used for intravenous drug delivery.

As intravenous injection is the most common method to administer therapeutics, it is critical to ensure that the nanoparticle reaches its targeted destination. Although research reported that encapsulated DNA into particles have modified their nanoparticles to be less toxic, have higher cellular uptake, or increase payload [41, 43, 74], it still have a limitation of off target delivery which causes systemic effects [75, 76]. For this reason, we investigated two endothelial cell specific antibodies, anti-EGFL7 and anti-Tie2+Tie1, on their ability to enhance cellular uptake in static and dynamic environments. We show that anti-Tie2+Tie1 has superior cellular uptake in both static and dynamic cell culture environments (**Figure 3 and Figure 5**). Compared to unconjugated nanoparticles, the targeting nanoparticles had significantly higher cellular uptake in the dynamic culture, supporting the notion that without targeting, the therapeutic may diffuse the efficacy of delivered.

In addition to site specific delivery, the encapsulated DNA needs to be bioactive. Others have shown that the sonication time or power, additives, or polymer molecular weight can affect the integrity of the plasmid [62]. We have shown that our synthesis method ensures plasmid delivery at several nanoparticle concentrations, and that the plasmid is bioactive. To find optimum concentration of NICD to upregulate Notch signaling related genes, the 100 and 250 μg nanoparticle of NICD encapsulating nanoparticles significantly upregulated Notch target genes, *Hes1 and Hay1,* compared to the gold standard lipofectamine with 5 μg of NICD (**Figure 4**). We show that GFP protein can be synthesized in HUVECs by delivering the plasmid. Additionally, Notch and its related genes were quantified showing upregulation. However, in 500 and 1,000 μg nanoparticle groups, while NICD was also significantly upregulated, expression levels of target genes were downregulated. This indicates that the 100 μg or 250 μg of NICD nanoparticle concentrations were preferred to induce a downstream genetic effect although the higher concentrations were able to increase expression of NICD. Accordance with previous report, increment of NICD does not proportionally increase target gene expression levels [28].

Although we have demonstrated PLGA nanoparticles with HMW (55–65 kDa) is appropriate material to deliver NICD plasmid to upregulated Notch signaling with *in vitro* flow experiment, we still need to test with *in vivo* experiment. Specifically, our *in vitro* experiment was limited in laminar flow while *in vivo* injection of nanoparticle would be exposed to pulsatile blood flow environment. In the future study, we will optimize the NICD plasmid concentration and conjugated with anti-Tie2+Tie1 coated PLGA nanoparticles for upregulated Notch signaling for animal model. This future experiment will help to translate our technology to effective therapeutic approach for translational medicine.

In this study, we have synthesized a PLGA nanoparticle that is capable of delivering plasmids to primary endothelial cells. In addition to being a nonviral transfection agent, the optimized nanoparticle was compatible with human cells and blood, and effectively delivered bioactive plasmid DNA to endothelial cells. These results demonstrate that PLGA targeting nanoparticles could increase the genetic delivery in complex environments, such as *in vivo*, with minimal adverse effects.

## Conclusion

In this work, we have shown that higher molecular weight PLGA outperforms the low molecular weight PLGA nanoparticles in cytotoxicity, cellular uptake, stability, and hemocompatibility. Additionally, the conjugation of anti-Tie2+Tie1 to the nanoparticles allows for a significant increase in endocytosis compared to those conjugated with anti-EGFL7. Lastly, our pCAG-GFP and TetO-FUW-NICD plasmids were both successfully encapsulated and transfected into HUVECs. Most importantly, the plasmid was bioactive after transfection as indicated by GFP imaging and RT-PCR analysis. In conclusion, we are able to show that plasmid loaded nanoparticles have a higher transfection efficiency, and create a significant genetic effect when applied to hard-to-transfect cells like HUVECs.

## Author Contributions

VM and AK: Nanoparticle characterization, hemocompatibility, antibody optimization; VM and UC: Plasmid loading, plasmid release analysis, 12 dyne/cm2 flow RT-PCR; VM and SL: Nanoparticle release study; VM, SL and LK: bacteria culture, plasmid isolation; JL and KN: conceptual design, research guide, data interpretation.

## Acknowledgements

Victoria Messerschmidt and Aneetta E. Kuriakose are supported by the National Institutes of Health (NIH) training award, T32 HL134613 (K.N). Juhyun Lee is supported by the American Heart Association 18CDA34110150 (J.L.) and NSF 1936519 (J.L.).

