## Supplementary document for "Notch Intracellular Domain Plasmid Delivery via Poly(lactic-co-glycolic acid) Nanoparticles to Upregulate Notch Signaling"

### Supplemental Data

Victoria L. Messerschmidt[1,2†], Aneetta E. Kuriakose[1,2†], Uday Chintapula[1], Samantha Laboy[1], Thuy Thi Dang Truong[1], LeNaiya A. Kydd[1], Justyn Jaworski[1], Kytai T. Nguyen[1,2*], Juhyun Lee[1,2*]

[1]Department of Bioengineering, University of Texas at Arlington, Arlington TX 76010 USA

[2]University of Texas Southwestern Medical Center, Dallas TX 75390 USA

[†] **These authors have contributed equally to this work**

### Supplementary Figures

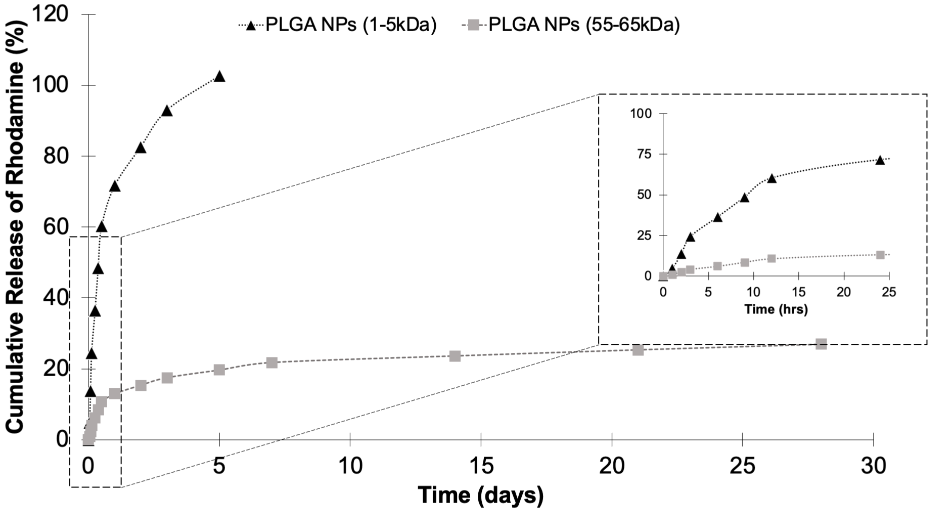


***Supplementary Figure 1: Rhodamine Release from High and Low Molecular Weight PLGA Nanoparticles.*** *Rhodamine Release in High and Low Molecular Weight PLGA. Release of Rhodamine into the supernatant has a burst release, followed by a sustained release up to 28 Days. The Low Molecular Weight nanoparticles release 100% of loaded rhodamine by 5 days. Inset shows initial burst release up to 24 hours.*

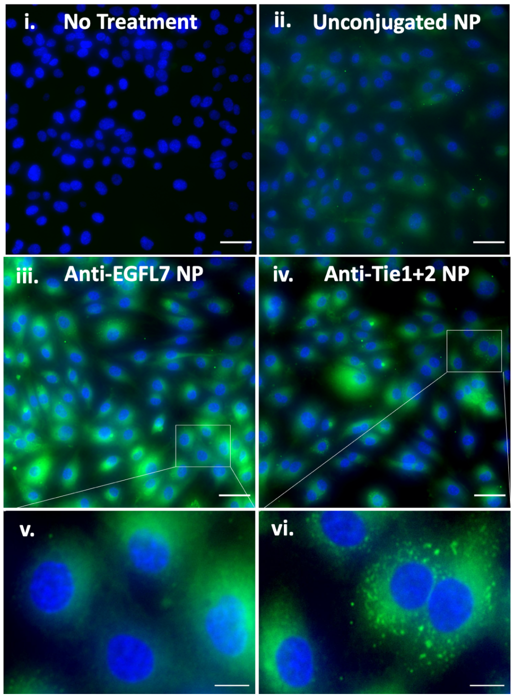


***Supplementary Figure 2: Static Culture of Antibody Conjugated Nanoparticles.*** *HUVEC's cultured with media only (i.), unconjugated nanoparticles (ii.), anti-EGFL7 conjugated nanoparticles (iii.), or anti-Tie2+Tie1 conjugated nanoparticles (iv.). Scale bar = 20 μm. Higher magnification of HUVEC's cultured in anti-EGFL7 conjugated nanoparticles (v.) or anti-Tie2+Tie1 conjugated nanoparticles. Scale bar = 5 μm.*

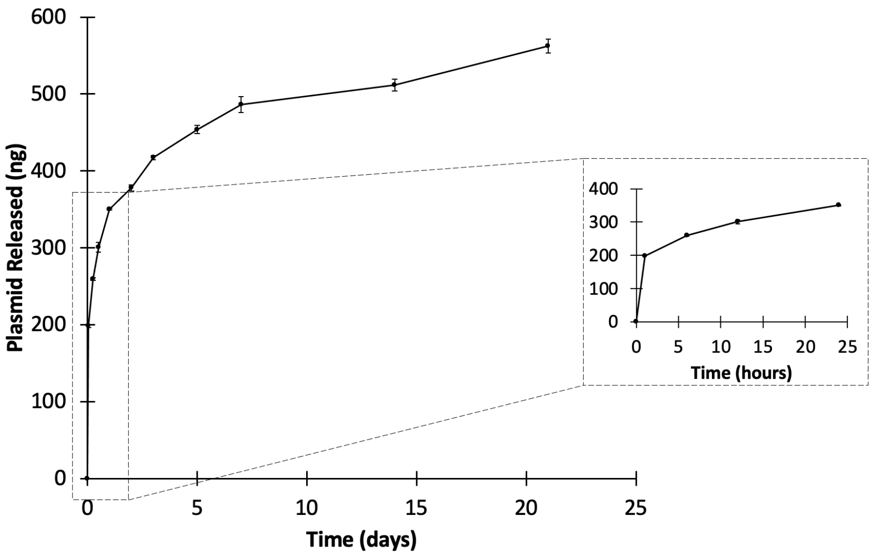


***Supplementary Figure 3: Characterization of GFP Plasmid-Loaded PLGA Nanoparticles.*** *GFP plasmid released from HMW PLGA nanoparticles over 21 days.*

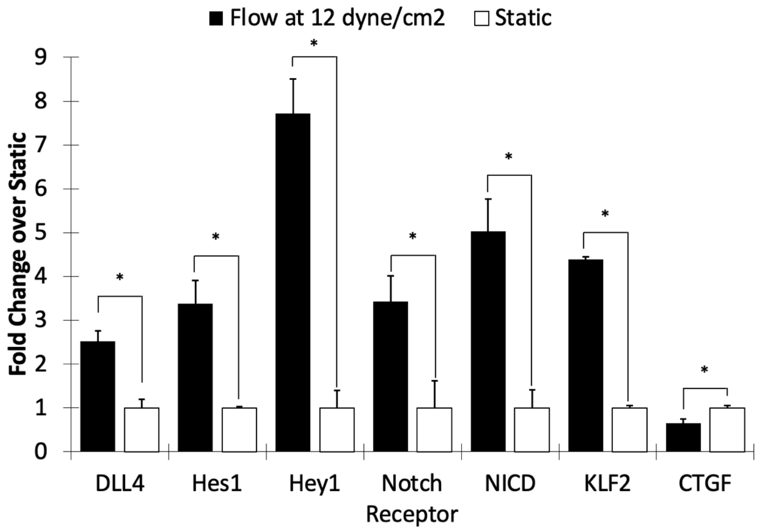


***Supplementary Figure 4: Natural Upregulation of Notch Related Genes due to Shear Stress.*** *RT-PCR results showing that notch related genes are upregulated from shear stress stimulus. KLF2 is also upregulated due to shear stress, while CTGF is downregulated.*

### Supplementary Tables

***Supplementary Table 1: Physical Characteristics of GFP Plasmid-Loaded PLGA Nanoparticles.***

| | **Size (nm)** | **Poly Dispersity** | **Zeta Potential (mV)** | **Encapsulation Efficiency (%)** |
|---|---|---|---|---|
| **GFP Plasmid-Loaded PLGA Nanoparticles** | 278.6 ± 47 | 0.22 ± 0.05 | -14.8 ± 2.0 | 38.9 ± 2.17 % |
